# Acyl-lipid desaturases and Vipp1 cooperate in cyanobacteria to produce novel omega-3 PUFA-containing glycolipids

**DOI:** 10.1101/2020.05.01.073155

**Authors:** Leslie B. Poole, Derek Parsonage, Susan Sergeant, Leslie R. Miller, Jingyun Lee, Cristina M. Furdui, Floyd H. Chilton

**Affiliations:** Department of Biochemistry, Wake Forest School of Medicine, Winston-Salem, NC; Center for Redox Biology and Medicine, Wake Forest School of Medicine, Winston-Salem, NC; Comprehensive Cancer Center, Wake Forest School of Medicine, Winston-Salem, NC; Department of Internal Medicine, Section on Molecular Medicine, Wake Forest School of Medicine, Winston-Salem, NC; Department of Physiology and Pharmacology, Wake Forest School of Medicine, Winston-Salem, NC; Department of Nutritional Sciences and the BIO5 Institute, University of Arizona, Tucson, AZ

## Abstract

**Background:** Dietary omega-3 (n-3), long chain (LC-, ≥ 20 carbons), polyunsaturated fatty acids (PUFAs) derived largely from marine animal sources protect against inflammatory processes and enhance brain development and function. With the depletion of natural stocks of marine animal sources and an increasing demand for n-3 LC-PUFAs, alternative, sustainable supplies are urgently needed. As a result, n-3 18 carbon and LC-PUFAs are being generated from plant or algal sources, either by engineering new biosynthetic pathways or by augmenting existing systems.

**Results:** We utilized an engineered plasmid encoding two cyanobacterial acyl-lipid desaturases (DesB and DesD, encoding Δ15 and Δ6 desaturases, respectively) and “vesicle-inducing protein in plastids” (Vipp1) to induce production of stearidonic acid (SDA,18:4 n-3) at high levels in three strains of cyanobacteria (10, 17 and 27% of total lipids in *Anabaena* sp. PCC7120, *Synechococcus* sp. PCC7002, and *Leptolyngbya* sp. strain BL0902, respectively). Lipidomic analysis revealed that in addition to SDA, the rare anti-inflammatory n-3 LC-PUFA eicosatetraenoic acid (ETA, 20:4 n-3) was synthesized in these engineered strains, and ∼99% of SDA and ETA was complexed to bioavailable monogalactosyldiacylglycerol (MGDG) and digalactosyldiacylglycerol (DGDG) species. Importantly, novel molecular species containing alpha-linolenic acid (ALA), SDA and/or ETA in both acyl positions of MGDG and DGDG were observed in the engineered *Leptolyngbya* and *Synechococcus* strains, suggesting that these could provide a rich source of anti-inflammatory molecules.

**Conclusions:** Overall, this technology utilizes solar energy, consumes carbon dioxide, and produces large amounts of nutritionally-important n-3 PUFAs and LC-PUFAs. Importantly, it can generate previously-undescribed, highly bioavailable, anti-inflammatory galactosyl lipids. This technology could therefore be transformative in protecting ocean fisheries and augmenting the nutritional quality of human and animal food products.

**Broader Context:** Dietary omega-3 (n-3), long chain polyunsaturated fatty acids (LC-PUFAs) typically found in marine products such as fish and krill oil are beneficial to human health. In addition to human consumption, most of the global supply of n-3 LC-PUFAs is used as dietary components for aquaculture. Marked increases in usage have created an intense demand for more sustainable, stable and bioavailable forms of n-3 PUFAs and LC-PUFAs. We utilized an engineered plasmid to dramatically enhance the production of 18-carbon and n-3 LC-PUFAs in three strains of autotrophic cyanobacteria. While the sustainable generation of highly valued and bioavailable nutritional lipid products is the primary goal, additional benefits include the generation of oxygen as a co-product with the consumption of only carbon dioxide as the carbon source and solar radiation as the energy source. This technology could be transformative in protecting ocean fisheries and augmenting the nutritional quality of human and animal food products. Additionally, these engineered cyanobacteria can generate previously undescribed, highly bioavailable, anti-inflammatory galactosyl lipids.

## Introduction

Eighteen-carbon (18C), omega-3 (n-3) polyunsaturated fatty acids (PUFAs) and particularly n-3 long chain (LC, ≥ 20 carbons) PUFAs have been shown to exert anti-inflammatory and cardioprotective roles in cardiovascular disease and several inflammatory diseases (1). Additionally, n-3 LC-PUFAs are essential for early childhood development, and deficiencies of n-3 LC-PUFAs are associated with mental disorders and cognitive decline (2-5). Consequently, several health organizations recommend increasing dietary consumption of n-3 PUFAs and LC-PUFAs, resulting in rapidly growing markets for these in functional foods, pharmaceuticals, dietary supplements and infant formulas (6, 7).

However, expansions in demand for n-3 PUFAs and n-3 LC-PUFAs have raised vital questions about their sustainability. For example, fish represent the predominant source of n-3 LC-PUFAs; however, wild caught fish are at or beyond exploitable limits, and more than half of fish consumed are farmed (7). Krill oil as another unsustainable source of n-3 LC PUFAs exerts even greater strains on the global health of ocean fisheries. Approximately 75% of the global supply of n-3 LC-PUFAs is currently utilized by aquaculture, which has led to a shift to n-6 PUFA-based vegetable (such as soybean and rapeseed) oil products, decreasing the nutritional quality of the farmed fish (8-11). Furthermore, there is a growing need for dietary n-3 PUFAs and n-3 LC-PUFAs in terrestrial livestock to enrich levels in meat, milk and egg products (12, 13).

Potential solutions to the growing demand for n-3 18C-PUFAs and LC-PUFAs are plant and algae-based sources produced through solar energy-dependent processes (7). Most plant-sourced, n-3 PUFA-containing oils, such as flaxseed oil, are enriched with the 18C-PUFA α-linolenic acid (ALA, 18:4, n-3), which has the potential to be converted to n-3 LC-PUFAs such as eicosapentaenoic acid (EPA, 20:5) and docosahexaenoic acid (DHA, 22:6). However, humans and most animals, including cold-water species of fish, are inefficient at converting ALA into EPA and DHA. The rate limiting steps in this conversion are the desaturation steps, and particularly the Δ6 desaturase (Fig. 1). However, the product of Δ6 desaturase, stearidonic acid (SDA, 18:4, n-3), bypasses this rate-limiting step; several human and animal studies show that seed oils containing SDA are more efficiently converted to EPA than those with ALA (14-17). SDA-containing seed oils from relatively rare plant species have been commercialized and common plant seed oils such as soybeans and canola have been genetically engineered to have enriched SDA content (20-29% of total fatty acids) (18, 19). Human clinical studies show that SDA-enhanced soybean oil significantly elevates n-3 LC-PUFAs and improves markers of cardiovascular health (20, 21). However, the feasibility of these commercial applications and the stability of these transgenic plants remain to be determined. More recently, there has been a marked increase in the production and sales in human consumer markets of n-3 LC-PUFAs (EPA and DHA) from phototrophic algae. Nevertheless, there are significant production cost barriers in supplying plant and animal sources of n-3 PUFAs and LC-PUFAs to the rapidly expanding aquaculture feed and livestock markets.

**Figure 1.**
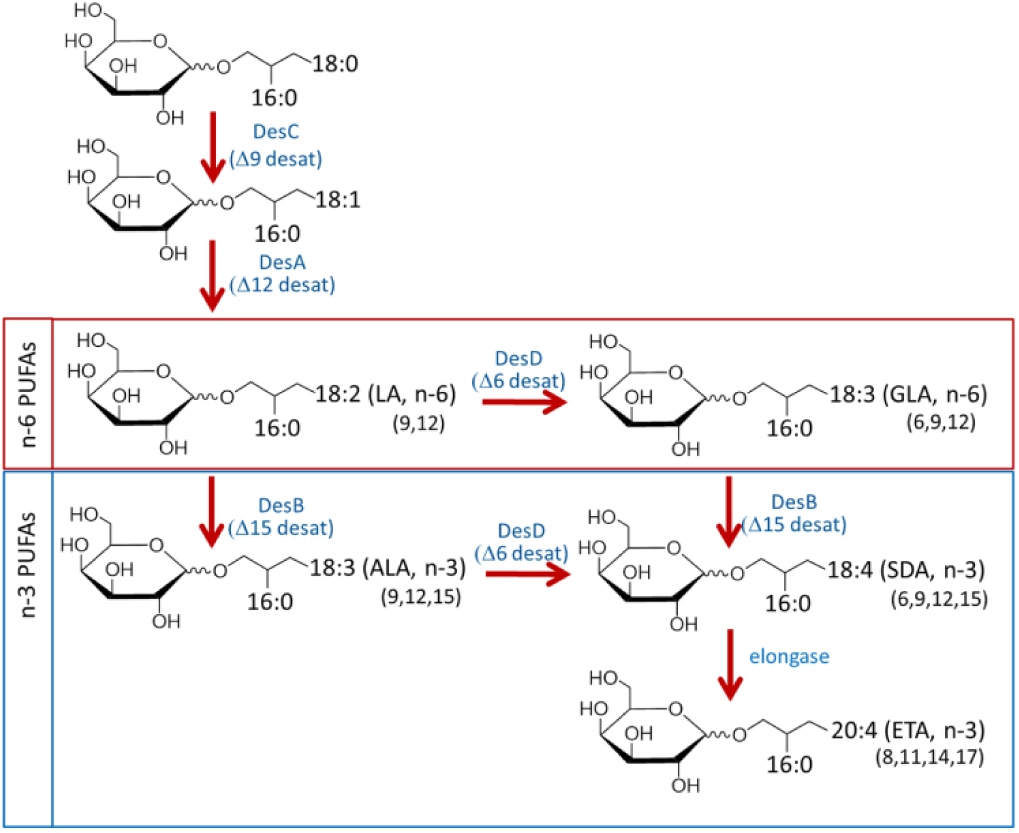
Cyanobacterial pathways of 18- and 20-carbon polyunsaturated fatty acid (PUFA) synthesis. Introduction of double bonds into stearic acid (18:0) involves a series of acyl-lipid desaturases designated DesC, DesA, DesD and DesB in cyanobacteria which catalyze desaturation at distinct sites of the carbon chain, ultimately producing stearidonic acid (SDA if all four desaturase steps occur. Addition of two more carbons by an elongase can then form eicosatetraenoic acid (ETA), the ω3 isomer of arachidonic acid. The major three n-3 (omega-3) polyunsaturated fatty acids observed in cyanobacteria are shown [alpha-linolenic acid (ALA), SDA and ETA]. The structures shown represent a monogalactosyldiacylglycerol (MGDG) backbone and typical 16:0 saturated fatty acid (palmitic acid) at the sn-2 position in addition to the unsaturated fatty acid at sn-1.

Cyanobacteria, which can be grown in large quantities requiring only sun, water and trace nutrients, have been subjected to mutagenesis and metabolic engineering for decades in pursuit of sustainable sources for numerous high-value products (22, 23). Genetic engineering of cyanobacteria through pathway modulation (both interruption or bolstering of existing pathways and introduction of new pathways) is enabling production of energy-containing molecules for use as biofuels (23-25). Efforts to augment lipid production through improved cyanobacterial hosts and pathway engineering have also been ongoing, but have met with challenges including low yields when the fatty acids produced are secreted into the media as free acids (26-28). In fact, PUFAs are naturally produced by cyanobacteria as essential constituents of the polar glycolipids in membranes, with the degree of unsaturation of membrane lipids controlling membrane fluidity (29, 30). The content of specific PUFAs varies among different cyanobacteria depending on the identity of the desaturase genes present in the genomes. Recent studies have suggested that these glycolipid-associated n-3 PUFAs are more bioavailable than fish and seed-based sources (31). Due to differences in digestive routes and physical forms (polar vs nonpolar lipids) of n-3 PUFA- and LC-PUFA-containing complex lipids, the bioavailability of diverse forms varies considerably. To date, most n-3 PUFAs or LC-PUFAs have been provided to humans and animals complexed to non-polar triglycerides from seed oils or marine fats. However, there are numerous problems with highly-enriched triglyceride formulations including the fact that large quantities of such concentrates are typically needed to achieve effective circulating and tissue (especially brain) levels of PUFAs and LC-PUFAs (32). To overcome these obstacles, ethyl esters, free fatty acids, re-esterified triglycerides or phospholipids (in the case of Krill oil) have been formulated, although with varying degrees of success. Cyanobacteria (and dark green plants with abundant chloroplasts) have thylakoid membranes that contain large quantities of the galactose-containing glycolipids monogalactosyldiacylglycerol (MGDG) and digalactosyldiacylglycerol (DGDG) (33, 34). Importantly, there is a selective pancreatic lipase (PLRP2) that mobilizes fatty acids from MGDG and DGDG (35), and initial rodent and human studies suggest that ingestion of LC-PUFAs with or complexed to these glycolipids improves their bioavailability (31, 36).

As the prokaryotic precursors of chloroplasts, cyanobacteria are biologically simpler than plants and algae, and genetic manipulation is generally more feasible, enabling metabolic reprogramming by engineering as noted above. Importantly for the purpose of this work, they also contain acyl-lipid desaturases and “Group 4” cyanobacteria have the critical four desaturases (DesC, DesA, DesB and DesD, Fig. 1) necessary to convert stearic acid (18:0) to SDA (37). However, as opposed to acyl-CoA or acyl-ACP desaturases, these desaturases act directly on fatty acids within the glycolipids, and these lipids account for ∼80% of total lipids in thylakoid membranes (38). With these understandings, the overall objective of the current study was to engineer cyanobacterial strains to augment the expression of three genes, *desB* and *desD* encoding acyl-lipid desaturases (known as Δ15 and Δ6 desaturases, respectively), and *vipp1* encoding a thylakoid membrane enhancing protein. The key question this work addresses is whether it is possible to markedly increase the capacity of cyanobacteria to produce SDA and eicosatetraenoic acid (ETA, 20:4, n-3) complexed to highly bioavailable MGDG and DGDG molecular species.

## Results

### Generation of plasmids and engineered cyanobacteria, and analyses of total fatty acid content

With the goal of maximizing cyanobacterial omega-3 production focused particularly on SDA, we selected three genes for overexpression which occur naturally in Group 3 and 4 cyanobacteria (37). Unlike the first two groups, which express just the Δ9 desaturase (Group 1), or both the Δ9 and Δ12 desaturases (DesC and DesA, respectively, Group 2), Group 3 and 4 cyanobacteria also express the Δ6 and/or Δ15 desaturases that enable production of the trienoic fatty acids GLA and ALA, and the tetraenoic fatty acid SDA (in the case of Group 4 cyanobacteria which have all four desaturases) (37). Specificities of the individual desaturases for the sites of double bond insertion in model cyanobacteria have been well established (39) and sequence signatures have emerged to facilitate functional assignment of new sequences (40, 41). While the Δ6 desaturase (DesD) acts on LA or ALA to generate GLA or SDA, respectively, the Δ15 desaturase (DesB) inserts a double bond three carbons from the methyl end, yielding omega-3 products ALA and SDA from LA and GLA, respectively (34, 42). We hypothesized that DesB and DesD overexpressed from a plasmid would impart or augment SDA synthesis in most cyanobacteria (i.e., those of Groups 2 through 4).

Previous studies reveal that these desaturase reactions occur within thylakoid membranes, and a thylakoid membrane formation enhancer gene, *vipp1* (which encodes Vesicle-inducing protein in plastids or Vipp1, also known as IM30) (38, 43), was the third protein selected for overexpression to potentially boost levels of newly-synthesized PUFAs formed by the enhanced desaturase system. All three synthetic genes (*desB, desD* and *vipp1*) encoding the authentic cyanobacterial protein products were incorporated into the expression plasmid pAM4418 (Fig. 2A) first described by Taton and colleagues (45), either singly or in combination with one or two other genes. The constructs were conjugated into *Leptolyngbya* sp. strain BL0902 (hereafter designated BL0902), a freshwater, filamentous cyanobacterium noted for its excellent growth characteristics and high lipid and especially LA content (44, 45). No obvious deleterious effects on growth were observed in any of the seven exconjugants.

**Figure 2.**
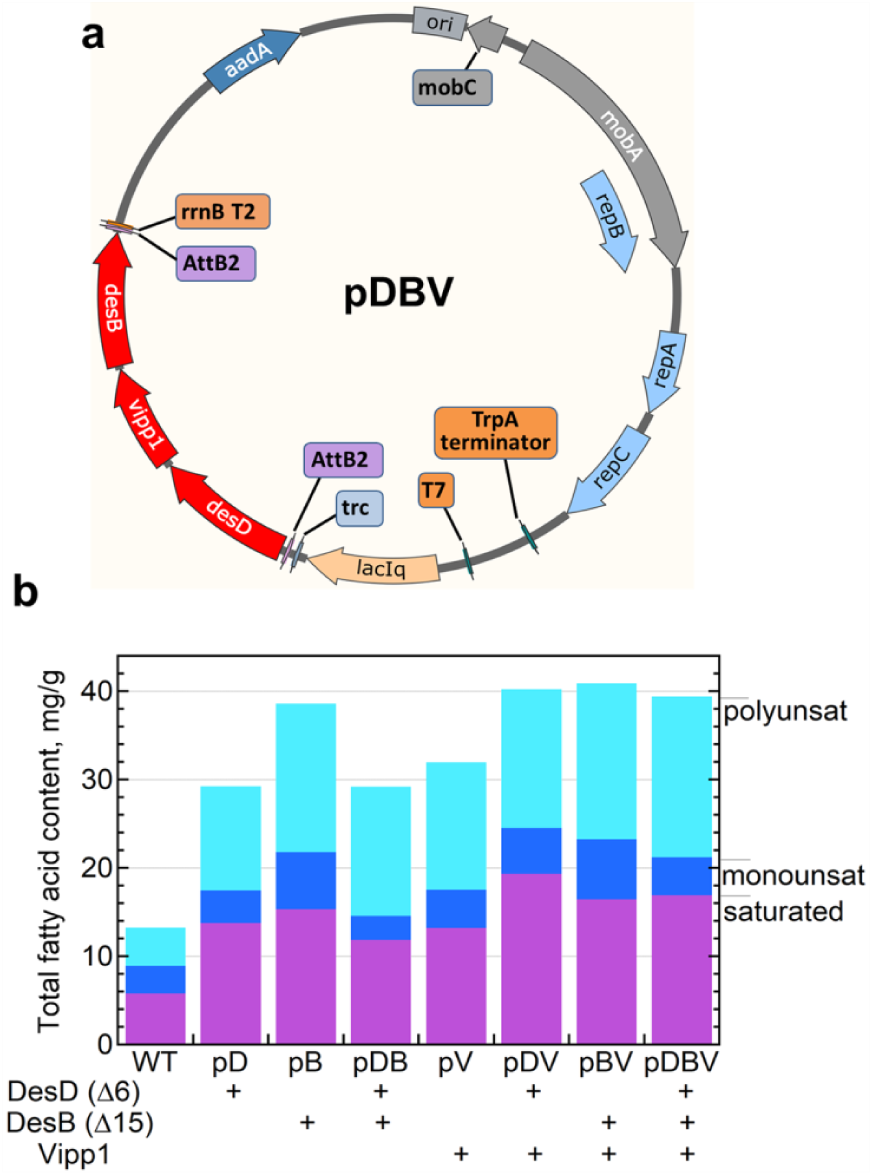
Map of the expression plasmid (pDBV), and fatty acid contents of constructs in *Leptolyngbya* BL0902. (A) To generate the pDBV (and other) plasmids, pAM4418-derived expression vectors were generated with synthetic genes designed to express: (i) the Δ6 desaturase (DesD) from *Synechocystis* sp. PCC 6803, (ii) the “Δ15” (ω3, or methyl-end) desaturase (DesB) from *Synechococcus* sp. PCC 7002, and/or (iii) the “vesicle-inducing protein in plastids” (Vipp1) from *Synechococcus* sp. PCC 7002. Included in the plasmid vector are *aad*A, conferring resistance to spectinomycin and streptomycin, as well as *trp*A and *rrn*B which block continued transcription. (B) Plasmids with one, two or all three of the inserted cyanobacterial genes were constructed and conjugated into *Leptolyngbya* sp. strain BL0902, transconjugants were selected on BG-11 agar plates containing spectinomycin and streptomycin, and cultures were grown at 30 °C in BG-11 media, harvested and dried for fatty acid analysis of lipid content by fatty acid methyl ester (FAME) analysis (gas chromatography with flame ionization detection, GC-FID). Shown are total saturated (magenta), monounsaturated (dark blue) and polyunsaturated (cyan) fatty acid contents (n=3 or more, from left to right, except n=2 for pDV).

All plasmid-bearing cells of BL0902 showed a marked elevation in both saturated and polyunsaturated fatty acids (with monounsaturated levels varying) compared with the wild type (Fig. 2B); the total fatty acid content of the seven exconjugants ranged from 2 to 2.5-fold greater after conjugation. The three gene-containing plasmid pDBV (Fig. 2A) increased the total fatty acid content from 13 to about 40 mg/g dry weight and total PUFAs from 5.8 to 16.9 mg/g dry weight (Fig. 2B and Table 1A). Addition of *vipp1* to all desaturase-expressing exconjugants elevated the total fatty acid content (Fig. 2B). To investigate the potential for the pDBV plasmid to impact SDA production in other cyanobacteria, we selected two strains for conjugation with our constructs, *Synechococcus* sp. PCC7002 and *Anabaena* sp. PCC7120 (subsequently referred to as 7002 and 7120). (*Synechococcus elongatus* PCC7942 was considered for engineering, but not further studied when initial fatty acid profiling indicated a lack of LA, see Supplementary Excel Data.) Unlike BL0902, strain 7002 engineered with pDBV showed no increase in total lipids while 7120 exhibited a modest increase of ∼ 23% (Fig. S1).

**Table 1.**
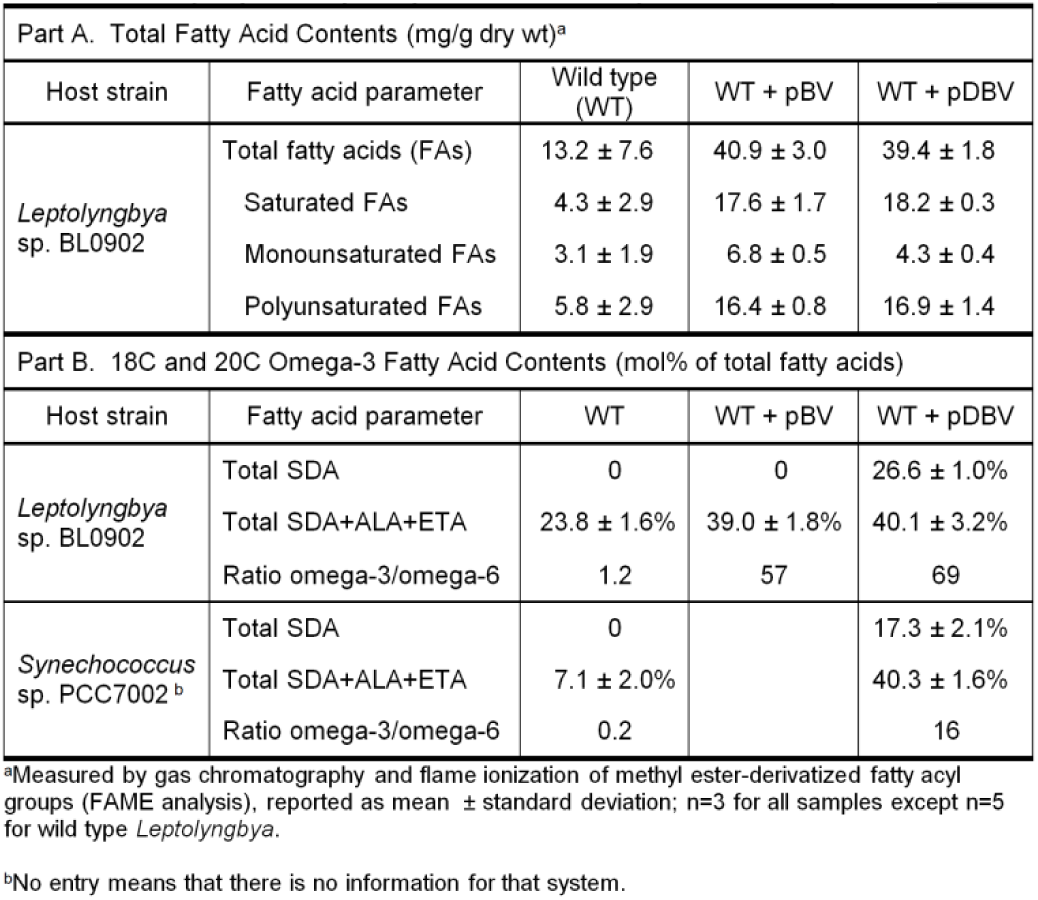
Summary of pBV and pDBV plasmid effects on cyanobacterial fatty acids.

### Modulation of individual fatty acids in engineered strains

Individual fatty acids (expressed as mg/g dry weight or percentage of total fatty acids) from the total lipid extracts of wild type and exconjugants of the three cyanobacterial strains were analyzed by gas chromatography-flame ionization detection (GC-FID) and mass levels compared (Supplementary Excel Data). Figure 3A illustrates the effect of inclusion of just one of the three genes on n-6 and n-3 PUFAs in BL0902, and Figure 3B summarizes the specific fatty acid contents when two or all three genes are included on the plasmid. Further information comparing the change in mass of pDBV-associated PUFAs with those from the BL0902 strains with the two-gene plasmids is also provided in the supplementary data (Fig. S2).

**Figure 3.**
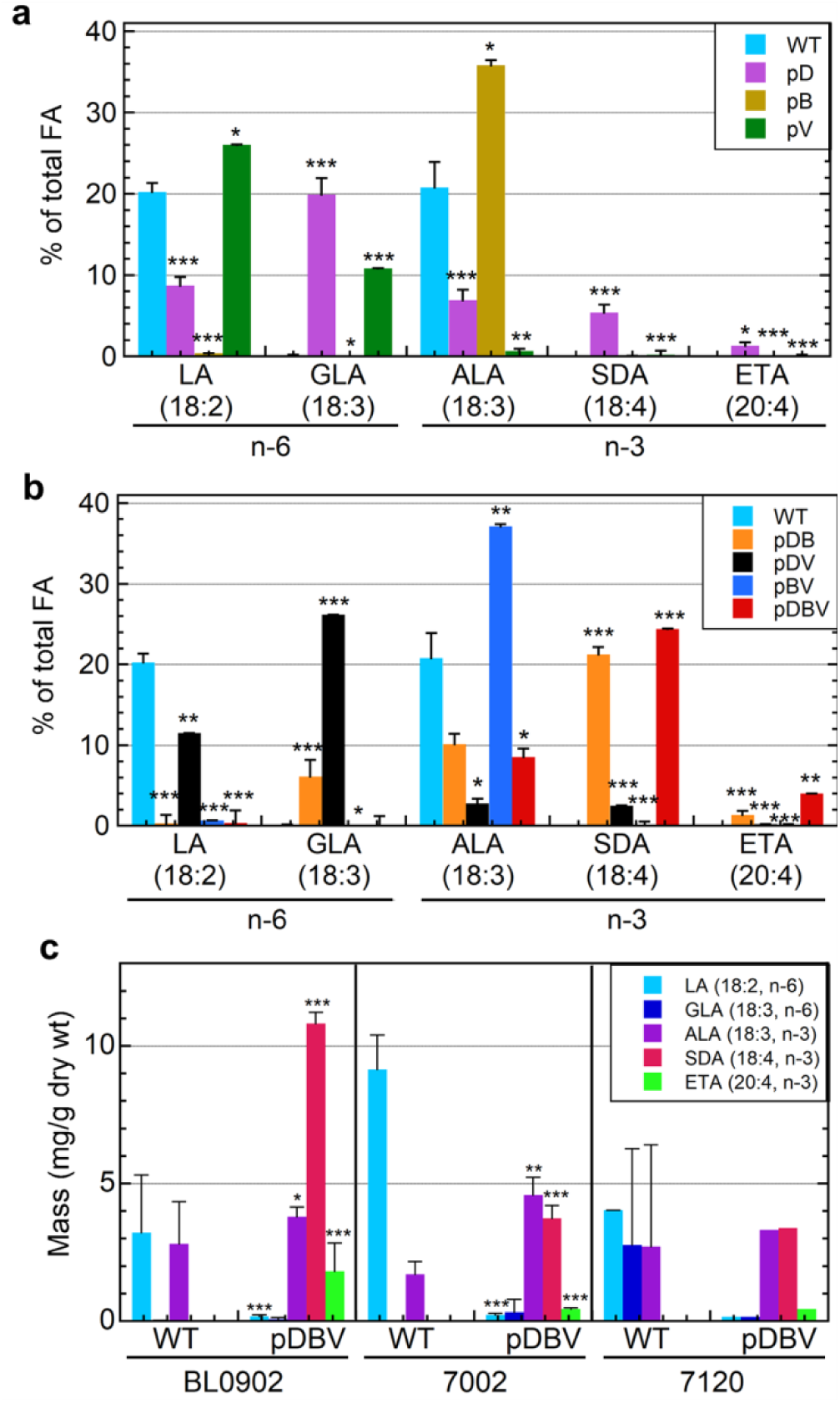
Quantitative analysis showing 18- and 20-carbon polyunsaturated fatty acids in wild type and engineered cyanobacteria. (A and B) PUFA analyses of plasmid-bearing *Leptolyngbya* sp. strain BL0902 as described in Figure 2B are shown as means ± standard deviation for wild type (WT) and single-gene constructs (A), or double and triple-gene constructs (B), expressed as the mol percent of total fatty acids (very similar to weight percent values). (C) PUFAs produced by WT and pDBV-bearing species of cyanobacteria, including *Leptolyngbya* sp. strain BL0902, *Synechococcus* sp. PCC 7002 and *Anabaena* sp. PCC 7120, are shown as averages of the mass (mg per g of dry weight) ± standard deviation (n=3 or more, from left to right, except n=2 and 1 for the 7120 samples). (A – C) Those exhibiting a statistically-significant difference in content compared with WT by a Students t-test are indicated with asterisks (*, p<0.05, **, p<0.01, ***, p<0.001).

In initial studies with wild type BL0902, we observed that only LA and ALA were present, implying that DesB (the Δ15 desaturase) is present and expressed to some extent (in addition to DesA and DesC), but the *des*D gene is likely missing (i.e., Leptolyngbya sp. strain BL0902 is likely a member of group 3α) (37). When only *des*D (encoding the Δ6 desaturase) is present on the plasmid, the organism now produces some amount of all five 18C and 20C PUFAs, with GLA predominating (consistent with the conversion of LA to GLA catalyzed by DesD). Maximal SDA is seen when both desaturases are included on the plasmid (Fig. 3B), particularly when *vipp*1 is also included (SDA as a percent of total FAs is not much changed between pDB and pDBV, but the total mass increases significantly with the latter, Fig. S2). With no *des*D included, expression of *des*B provided by the plasmid has a strong effect and leads to considerable accumulation of ALA at the expense of LA in BL0902 with constructs pB and pBV (gold in Fig. 3A and dark blue in Fig. 3B, Table 1) consistent with the Δ15 desaturase activity of DesB (Fig. 1). Importantly, the full three-gene plasmid shifted the profile to the greatest quantities of SDA and ETA, with some ALA remaining but no LA or GLA (red in Figs. 3B and 3C). Total SDA contents reached 26.6 ± 1.0 mol% of total fatty acids in the pDBV exconjugant, and the total n-3 PUFAs including LC-PUFAs (ALA, SDA and ETA) reached 40% of the lipid content in BL0902 (Table 1). The remarkable shift from n-6 to n-3 PUFA-containing lipids in the pDBV exconjugant was limited from producing additional SDA and ETA primarily by the unavailability of precursor substrates, LA and GLA. With such a depletion of LA and GLA, the pDBV exconjugant produced a ratio of n-3 to n-6 PUFAs of 69:1 (Table 1).

The two other strains of cyanobacteria (7002 and 7120) were tested with and without pDBV and like BL0902, wild type strains contained no detectable SDA or ETA, but both were produced upon addition of pDBV (Fig. 3C). Quantities of SDA produced by 7002 and 7120 were 31-35% of that generated by the engineered BL0902 strain (Fig. 3C), an organism which was noted for its favorable lipid composition when first isolated (44). Although not depicted in Figure 3, small amounts of the n-3 LC-PUFAs EPA (20:5, n-3) and docosapentaenoic acid (DPA; 22:5, n-3) were also observed in some of the engineered (but not wild type) strains (Supplementary Excel Data). Mass quantities of 0.03 ± 0.02 and 0.19 ± 0.05 mg/g dry weight were obtained for EPA in engineered 7002 and BL0902, respectively, while DPA was observed at 0.25 mg/g dry weight in only 7120 (Fig. S1).

### Lipidomics analysis

Lipidomics analysis was performed by liquid chromatography with tandem mass spectrometry (LC-MS/MS) using a high resolution Q Exactive HF Hybrid Quadrupole - Orbitrap mass spectrometer and was utilized to determine the molecular classes and species of all lipids, but especially those containing ALA, SDA and ETA. More than 300 lipid molecular species were identified in wild type BL0902 (309 total) and 258 species in the pDBV exconjugants (n=4 per group). Figure 4 illustrates that there are 159 lipid molecular species shared by the wild type and pDBV exconjugants, while 99 lipid molecular species were unique to the exconjugants. These unique lipids included ALA-, SDA-, ETA-containing molecular species of MGDG and DGDG classes (Fig. 4B and 4C). Importantly, SDA is found almost entirely in the MGDG and DGDG species (Table 2, Figs. 4B and 4C). In contrast, ALA is predominantly in MGDG and DGDG in WT, but is highly enriched in phosphatidyl glycerol (PG, 56%) in the pDBV exconjugant. ETA is found predominantly in MGDG (84%) with none detected in DGDG (Fig. 4B).

**Figure 4.**
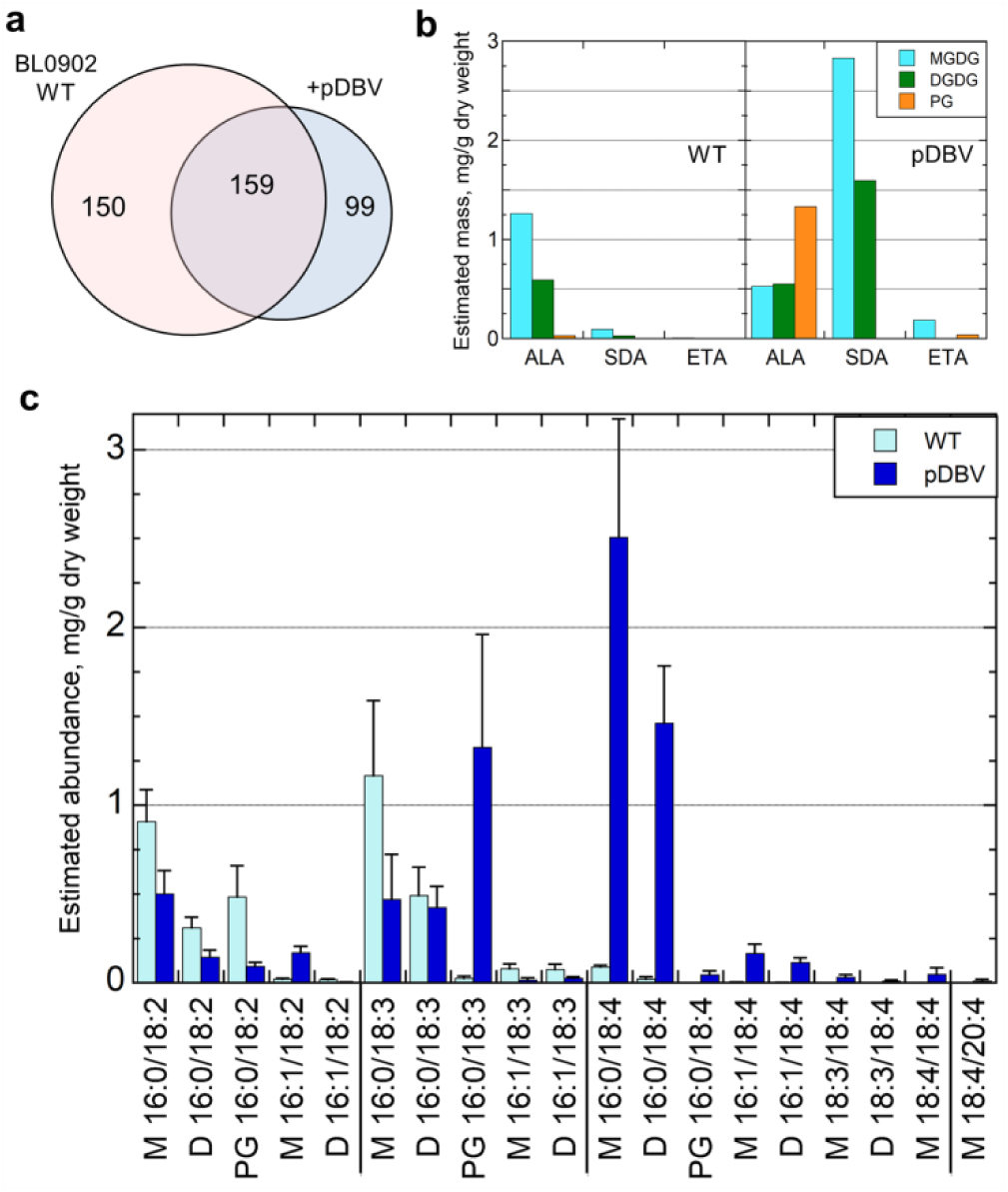
Lipid molecular species by LC-MS/MS of PUFA distribution in engineered and wild type *Leptolyngbya* BL0902. LC-MS/MS was conducted on lipid extracts dissolved in isopropyl alcohol/methanol (50:50), chromatographed on an Accucore C30 column, and introduced by heated electrospray ionization into a Q Exactive HF Hybrid Quadrupole - Orbitrap Mass Spectrometer with MS scans collected in data dependent mode, as described in Methods. (A) Venn diagram of the number of distinct molecular species observed for *Leptolyngbya* BL0902 without (WT) or with conjugation with pDBV (n=4 for each). (B and C) 18- and 20-carbon PUFAs were observed to be complexed to monogalactosyldiacylglycerol (MGDG), digalactosyldiacylglycerol (DGDG), or phosphatidyl glycerol (PG). (B) Distribution of ALA (18:3, n-3), SDA (18:4, n-3) and ETA (20:4, n-3) among the three glycolipids is shown, illustrating the selectivity for the lipid backbone for each PUFA. (C) Shown in light blue (WT) and dark blue (engineered with pDBV) are the mean ± standard deviation of estimates of mg/g of fatty acids based on (i) normalized peak areas from LC/MS, (ii) fraction of total peak area for each species in a sample, and (iii) known total fatty acid yield for that organism from GC-FID analysis. This treatment assumes that all species exhibit the same ionization efficiency. Species across the bottom refer to the two acyl chains associated with MGDG (M), DGDG (D) or PG. Note the shift from fewer to more double bonds upon introduction of the pDBV plasmid.

**Table 2.** Lipidomics analysis of molecular species in wild type (WT) and engineered (+pDBV) *Letolyngbya*

| Molecular species<sup>b</sup> | MGDG<sup>c</sup> |  | DGDG |  | SQDG |  | PG |  |
| --- | --- | --- | --- | --- | --- | --- | --- | --- |
|  | WT | +pDBV | WT | +pDBV | WT | +pDBV | WT | +pDBV |
| 14:0/16:0 |  |  | 0.005±0.001 | 0.002±0.001 |  |  | 0.001±0.001 | 0.000±0.000 |
| 14:0/18:3 |  |  | 0.013±0.001 | 0.010±0.003 |  |  |  |  |
| 14:0/18:4 | <u>0.000±0.000</u> | <u>0.009±0.002</u> | <u>0.000±0.000</u> | <u>0.006±0.002</u> |  |  |  |  |
| 16:0/16:0 | 0.066±0.024 | 0.129±0.038 | 0.043±0.022 | 0.150±0.111 | 0.008±0.005 | 0.016±0.015 | 0.037±0.023 | 0.113±0.037 |
| 16:0/16:1 | 0.161±0.028 | 0.201±0.080 | 0.045±0.018 | 0.078±0.024 |  |  | 0.077±0.025 | 0.089±0.028 |
| 16:0/16:2 | 0.021±0.006 | 0.016±0.005 | 0.011±0.004 | 0.006±0.001 |  |  | 0.004±0.002 | 0.007±0.003 |
| 16:0/16:3 | 0.001±0.000 | 0.040±0.018 |  |  |  |  |  |  |
| 16:0/16:4 | 0.000±0.000 | 0.010±0.003 |  |  |  |  |  |  |
| 16:0/17:0 | 0.002±0.001 | 0.007±0.002 | 0.002±0.002 | 0.012±0.005 |  |  | 0.000±0.001 | 0.001±0.002 |
| 16:0/17:1 | 0.040±0.010 | 0.066±0.020 | 0.009±0.004 | 0.016±0.003 |  |  | 0.013±0.005 | 0.044±0.020 |
| 16:0/17:2 | 0.028±0.006 | 0.034±0.012 | 0.011±0.003 | 0.009±0.002 |  |  | 0.005±0.002 | 0.001±0.002 |
| 16:0/17:3 | 0.020±0.008 | 0.024±0.004 | 0.010±0.006 | 0.010±0.006 |  |  | 0.000±0.000 | 0.014±0.005 |
| 16:0/17:4 | 0.000±0.000 | 0.045±0.014 | 0.000±0.000 | 0.015±0.003 |  |  |  |  |
| 16:0/18:0 | 0.008±0.005 | 0.020±0.019 | 0.007±0.008 | 0.029±0.011 |  |  | 0.002±0.002 | 0.006±0.002 |
| 16:0/18:1 | 0.410±0.088 | 0.488±0.180 | 0.087±0.044 | 0.185±0.057 |  |  | 0.275±0.068 | 0.397±0.176 |
| 16:0/18:2 | 0.908±0.179 | 0.501±0.132 | 0.311±0.059 | 0.145±0.039 |  |  | 0.484±0.177 | 0.093±0.022 |
| 16:0/18:3 | 1.166±0.421 | 0.469±0.254 | 0.491±0.160 | 0.424±0.118 |  |  | 0.027±0.013 | 1.325±0.636 |
| 16:0/18:4 | <u>0.090±0.011</u> | <u>2.507±0.665</u> | <u>0.022±0.013</u> | <u>1.462±0.321</u> |  |  | <u>0.001±0.001</u> | <u>0.046±0.021</u> |
| 16:0/19:1 | 0.007±0.001 | 0.017±0.006 | 0.002±0.001 | 0.004±0.001 |  |  |  |  |
| 16:0/19:2 | 0.006±0.004 | 0.002±0.001 | 0.003±0.002 | 0.001±0.000 |  |  |  |  |
| 16:0/19:3 | 0.002±0.001 | 0.007±0.001 |  |  |  |  |  |  |
| 16:0/20:0 | 0.000±0.000 | 0.001±0.001 |  |  |  |  |  |  |
| 16:0/20:2 | 0.004±0.003 | 0.003±0.001 | 0.003±0.003 | 0.001±0.000 |  |  |  |  |
| 16:0/20:3 | 0.012±0.009 | 0.033±0.010 | 0.005±0.004 | 0.010±0.003 |  |  | 0.001±0.002 | 0.004±0.001 |
| 16:0/20:4 | <u>0.006±0.004</u> | <u>0.174±0.065</u> |  |  |  |  | <u>0.000±0.000</u> | <u>0.037±0.021</u> |
| 16:0/20:5 | <u>0.077±0.010</u> | <u>0.122±0.070</u> |  |  |  |  |  |  |
| 16:1/16:1 | 0.005±0.001 | 0.007±0.002 | 0.005±0.001 | 0.003±0.001 |  |  |  |  |
| 16:1/16:2 | 0.001±0.001 | 0.000±0.000 |  |  |  |  |  |  |
| 16:1/17:3 | 0.001±0.001 | 0.001±0.000 | 0.001±0.001 | 0.000±0.000 |  |  |  |  |
| 16:1/18:2 | 0.022±0.004 | 0.171±0.036 | 0.019±0.005 | 0.004±0.001 |  |  |  |  |
| 16:1/18:3 | 0.080±0.028 | 0.018±0.011 | 0.076±0.030 | 0.026±0.009 |  |  | 0.000±0.000 | 0.006±0.001 |
| 16:1/18:4 | <u>0.003±0.001</u> | <u>0.166±0.051</u> | <u>0.002±0.001</u> | <u>0.115±0.027</u> |  |  |  |  |
| 17:0/18:3 | 0.001±0.000 | 0.002±0.001 |  |  |  |  |  |  |
| 17:0/18:4 | <u>0.001±0.000</u> | <u>0.006±0.004</u> |  |  |  |  |  |  |
| 17:1/18:3 | 0.001±0.001 | 0.001±0.001 | 0.001±0.001 | 0.000±0.000 |  |  |  |  |
| 17:5/23:2 |  |  |  |  | 1.089±0.123 | 2.172 ±0.449 |  |  |
| 17:5/23:3 |  |  |  |  | 0.005±0.003 | 0.006±0.003 |  |  |
| 18:0/18:1 | 0.001±0.001 | 0.002±0.001 |  |  |  |  | 0.001±0.000 | 0.002±0.001 |
| 18:1/18:2 | 0.005±0.005 | 0.001±0.001 |  |  |  |  | 0.001±0.000 | 0.008±0.003 |
| 18:1/18:3 |  |  | 0.002±0.001 | 0.071±0.027 |  |  |  |  |
| 18:2/18:2 | 0.009±0.003 | 0.011±0.004 | 0.003±0.002 | 0.001±0.001 |  |  |  |  |
| 18:2/18:3 | 0.005±0.004 | 0.005±0.004 | 0.005±0.003 | 0.006±0.005 |  |  |  |  |
| 18:3/18:3 | 0.003±0.003 | 0.001±0.001 | 0.002±0.001 | 0.000±0.000 |  |  |  |  |
| 18:3/18:4 | <u>0.000±0.000</u> | <u>0.032±0.015</u> | <u>0.000±0.000</u> | <u>0.012±0.004</u> |  |  |  |  |
| 18:4/18:4 | <u>0.000±0.000</u> | <u>0.049±0.037</u> |  |  |  |  |  |  |
| 18:4/20:4 | <u>0.000±0.000</u> | <u>0.012±0.009</u> |  |  |  |  |  |  |

<sup>a</sup>Values reported are estimates of mg/g of total fatty acid based on (i) normalized peak areas from LC/MS, (ii) fraction of total peak area for each species in a sample, and (iii) known total fatty acid yield for that organism from GC-FID analysis. This treatment assumes that all species exhibit the same ionization efficiency. Shown are mean +/- standard deviation for wild type (WT) *Leptolyngbya* BL0902 and the same strain with pDBV (n=4 for each). Molecular species containing at least one acyl group with SDA (18:4), ETA (20:4) or EPA (20:5) are underlined. No entry means that the species was not observed in the WT or pDBV samples in that category.
<sup>b</sup>Molecular species shown are for the two fatty acyl chains, giving carbon chain length and number of double bonds for each (e.g., 16:0 means 16 carbons long with 0 double bonds).

The individual molecular species of MGDG, DGDG and PG are illustrated in Figure 4C. Glycolipids contain 18C acyl groups such as 18:0, 18:1 (n-9), 18:2 (n-6), 18:3 (n-3 or n-6) at the *sn*-1 position and C16 acyl groups including 16:0, 16:1, 16:2, 16:3 at the *sn*-2 position of the molecule (34). In the case of MGDG, DGDG, and PG containing ALA, SDA or ETA, the most common fatty acid observed at the *sn-*2 acyl position is palmitic acid (16:0) (89-95% of the time, Fig. S3). Interestingly, the pDBV exconjugants produced several novel and unexpected MGDG and DGDG molecular species containing 18C and/or 20C PUFAs in both (*sn*-1 and *sn*-2) positions, including SDA and ALA, SDA and SDA, or SDA and ETA (Fig. 4C).

A second experiment comparing BL0902 and 7002 samples with and without pDBV demonstrated similar molecular profiles, with ALA/ALA, ALA/SDA, ALA/ETA, SDA/SDA and SDA/ETA “double omega-3” chains among the MGDG and DGDG molecular species, as well as the SDA/SDA molecular species in sulfoquinovosyldiacylglycerol (SQDG). In fact, the main difference between the outputs from the two experiments was an accumulation of SDA and especially ALA in SQDG rather than PG (Table S2). There also were other rarer PUFAs induced by pDBV in both BL0902 and 7002 including 16:3, 16:4, 17:2, 17:4, 20:3, and 20:5 (Supplementary Excel Data and Table S1). Another highly unusual fatty acid, 18:5, was present in MGDG and SQDG in the second experiment and the SQDG in particular was induced upon addition of pDBV to both BL0902 and 7002.

## Discussion

The primary objective of this work was to determine the capacity of cyanobacteria strains to produce SDA as well as elongation and desaturation metabolites (ETA and EPA, respectively) complexed to potentially more bioavailable glycolipids (MGDG and DGDG). This was accomplished by conjugal transfer into three cyanobacteria of a three-gene plasmid, pDBV, which encodes the thylakoid membrane-promoting protein Vipp1 and two acyl-lipid desaturases (Δ6 and Δ15 desaturases) that occur naturally in cyanobacteria. The total yield of fatty acids increased three-fold in the pDBV exconjugants, and SDA levels which were at baseline in the wild type cyanobacteria rose to 26.6 mol% in *Leptolyngbya* BL0902 and 17.3 mol% in *Synechococcus* PCC 7002. Importantly, n-3 PUFAs and LC-PUFAs (ALA+SDA+ETA) comprised ∼40% of total fatty acids in engineered *Leptolyngbya* BL0902, and these were incorporated into MGDG and DGDG with a n-3 to n-6 PUFA ratio of >50:1 (Table 1). In comparison, similar studies reported by Chen et al. (46) expressing tagged versions of DesB, or both DesB and DesD, in *Synechocystis* sp. PCC6803 achieved levels of SDA of around 10.8% and 13.1%, respectively (less than half as much as we observed with pDBV-engineered *Leptolyngbya* BL0902) when grown at 20 °C, less if grown at the higher temperature of 30 °C that we used here (Table 3). In terms of total mass accumulated, SDA in our pDBV *Leptolyngbya* was produced (in about mid-log phase) at 10.8 ± 0.4 mg/g dry weight, which is comparable to the maximum observed previously of 12.2 ± 2.4 mg/g by Yoshino et al. under high incident light [150 µmol photons/(m^2^ · s), compared with 20-30 µmol photons/(m^2^ · s) in our studies] (Table 3) (47).

**Table 3.** Content of 18C omega-3 fatty acids in engineered cynobacteria from this and previous studies.

| Species | Vector for engineering | Temp (°C) | Photon flux density, $\mu\text{mol photons (m}^2 \cdot \text{s)}^{-1}$ | ALA (18:3, n-3), mol% of total FAs<sup>a</sup> | SDA (18:4, n-3), mol% of total FAs<sup>a</sup> | Reference |
| --- | --- | --- | --- | --- | --- | --- |
| *Leptolyngbya* sp. strain BL0902 | None (wt) | 30 | 20-30 | 22.6 ± 11.5 | n.d.<sup>b</sup> | This study |
| | With pBV (to express DesB and Vipp1)<sup>c</sup> | 30 | 20-30 | 37.7 ± 1.7 | n.d. | |
| | With pDBV (to express DesD, DesB and Vipp1)<sup>c</sup> | 30 | 20-30 | 9.3 ± 0.9 | 26.6 ± 1.0 | |
| *Synechococcus elongatus* PCC 7942 | None (wt) | 22 | 60 | n.d. | n.r.<sup>d</sup> | Santos-Merino et al., 2018 |
| | MSM26 (strain with overexpressed *desA* and *desB*)<sup>e</sup> | 22 | 60 | 7.6 ± 1.5 | n.r. | |
| | | 22 | 250 | 8.8 ± 1.8 | n.r. | |
| | MSM45 (strain with $\Delta fabD$ (deleted), and overexpressed *fabF*, *desA* and *desB*)<sup>e</sup> | 22 | 60 | 22.6 ± 1.5 | n.r. | |
| *Synechococcus* sp. strain NKBG 15041c | Empty vector | 25 | 50 | ~21 ± 4<sup>f</sup> | n.d. | Yoshino et al., 2017 |
| | Vector for *desD* overexpression<sup>g</sup> | 25 | 50 | 20.6 ± 3.7 | 2.1 ± 0.5 | |
| | | 25 | 100 | 45.7 ± 7.4 | 5.1 ± 0.6 | |
| | | 25 | 150 | 53.1 ± 1.3 | 6.2 ± 0.5 | |
| | | 30 | 50 | 9.0 ± 6.3 | 0.9 ± 0.4 | |
| *Synechococcus* sp. PCC7002 | None (wt) | 30 | 60 | 5.2 ± 0.3 | n.d. | Dong et al., 2015 |
| | Syd15D (for *desB* overexpression)<sup>h</sup> | 30 | 60 | 28.3 ± 2.3 | n.d. | |
| | Syd6Dd15D (for *desD* and *desB* overexpression)<sup>g,h</sup> | 30 | 60 | 6.6 ± 0.7 | 11.6 ± 1.6 | |
| *Synechocystis* sp. PCC 6803 | None (wt) | 20 | 40 | 2.5 ± 0.3 | 1.5 ± 0.2 | Chen et al., 2014 |
| | | 30 | 40 | 1.5 ± 0.2 | 1.2 ± 0.3 | |
| | Sy15 (for *desB* overexpression)<sup>i</sup> | 20 | 40 | 23.1 ± 2.3 | 10.8 ± 1.6 | |
| | | 30 | 40 | 17.5 ± 2.3 | 9.1 ± 1.3 | |
| | Sy15Sy6 (for *desB* and *desD* overexpression)<sup>i,j</sup> | 20 | 40 | 16.4 ± 1.9 | 13.1 ± 1.3 | |
| | | 30 | 40 | 23.6 ± 3.4 | 7.8 ± 0.7 | |

<sup>a</sup> Expressed as mean ± standard deviation. <sup>b</sup> n.d. = not detected. <sup>c</sup> pBV encodes *Synechococcus* sp. PCC7002 DesB for $\Delta 15$ desaturase (also known as $\omega 3$ or methyl end desaturase) and *Synechococcus* sp. PCC7002 Vipp1 for inducing thylakoid membranes; pDBV is the three-gene plasmid derived from pAM4418 (Fig. 2) developed in this work which includes the two genes of pBV plus *Synechocystis* sp. PCC6803 *desD* for $\Delta 6$ desaturase expression. <sup>d</sup> n.r. = not reported. <sup>e</sup> *desA* and *desB* were from *Synechococcus* sp. PCC7002. <sup>f</sup> Estimated from mass if total fatty acid content is same as transformed strain. <sup>g</sup> *desD* was from *Synechocystis* sp. PCC6803. <sup>h</sup> *desB* was from *Synechocystis* sp. PCC6803. <sup>i</sup> *desB* was from *Synechocystis* sp. PCC6803 plus a C-terminal FLAG tag.

The rationale for the addition of *vipp1* to the plasmid was to enhance thylakoid membranes and thus the content of MGDG and DGDG as well as the potential activities of the acyl-lipid desaturases (DesC, DesA, DesB and DesD). While Vipp1 is reported to promote thylakoid membrane biogenesis and maintenance in plants and cyanobacteria associated with enhanced photosynthetic machinery (48, 49), the effect of this gene on PUFA and LC-PUFA content has not been previously reported. It is well established that n-3 PUFA contents of cyanobacteria are significantly affected by temperature and light intensity (47, 50-52). The SDA and ETA contents may therefore be even further enhanced in these cyanobacteria by growth at lower temperatures and higher light intensities and further engineering of the cyanobacteria to enhance their capacity to generate more precursor substrates (LA, GLA or ALA) necessary for SDA and ETA production. For example, *Spirulina* strains can contain ∼50% of their total fatty acids as LA and GLA (53), and other cyanobacterial strains have been engineered that contain 25-82% of their total fatty acids as LA and ALA (46, 47, 54).

Given SDA’s stability relative to n-3 LC-PUFAs in food matrices and its capacity to be more efficiently (than ALA) converted to health-promoting EPA in humans, fish and livestock, a great deal of effort has gone into finding natural systems and designing new engineered pathways that produce high quantities of SDA and high ratios of n-3 to n-6 PUFAs (55). SDA is seldom naturally found in cyanobacteria (56) unless at least one acyl-lipid desaturase is provided on a multicopy vector (47), and even then, the maximal content of 26.6% SDA reached in the current study is as we have noted at least two-fold higher than previous engineered strains (Table 3) (46, 47). SDA-producing transgenic soybean oil contains 20-26% of the total fatty acids as SDA with n-3 to n-6 PUFA ratios of ∼ 1 or lower (19, 21, 55, 57), and seed oil from *Echium plantagineum* naturally contains ∼13% of the total fatty acids as SDA (15, 58, 59).

Lipidomics analysis identified complex lipids and individual molecular species containing newly-synthesized n-3 PUFAs and LC-PUFAs. Greater than 99% of SDA and ETA resided in MGDG and DGDG with the majority at the *sn*-1 position and palmitic acid at the *sn*-2 position of the glycolipid backbone (Fig. 4B and Fig. S3). Additionally, several highly unusual MGDG and DGDG molecular species containing SDA at both acyl positions or ALA:SDA, ALA:ETA and SDA:ETA combinations were detected. ETA is a rare n-3 LC-PUFA in nature comprising ∼ 1% of fish oils. ETA is also found in triglycerides of transgenic seeds from *Camelina* and in New Zealand green-lipped mussel (*Perna canaliculus*) (60, 61). ETA (also known as omega-3 arachidonic acid), an elongation product of SDA, is a structural analog of the n-6 arachidonic acid. Previous studies have demonstrated ETA’s capacity to serve as a dual inhibitor of cyclooxygenases (COX1 and COX2) and lipoxygenases that can block the production of several classes of pro-inflammatory eicosanoids including leukotrienes, prostaglandins, and thromboxanes (62, 63). ETA has also been demonstrated to compete with arachidonic acid at the arachidonoyl-CoA synthetase step thereby preventing arachidonic acid uptake (61, 64). Importantly, lipid extracts from New Zealand green-lipped mussel also have been shown to have benefits in patients with atopic asthma (65). Future studies will determine whether molecular species of MGDG and DGDG such as ALA:ETA, SDA:ETA and ETA:ETA have the potential to serve as bioavailable anti-inflammatory compounds.

Significant advantages of sourcing SDA and ETA from cyanobacteria include: 1) they require minimal nutritional demands, relying on photosynthesis and carbon dioxide rather than fermentable sugars, and do not require arable land for production (66); 2) they are able to serve directly as single ingredient feeds for aquaculture and livestock (simply by drying and being fed as flakes or pellets), therefore offering the possibility of less labor- and land-intensive cultivation of such feeds (22); 3) cyanobacteria such as *Spirulina* are currently used as food supplements because of their high protein content and digestibility (22); and 4) SDA and ETA formed in cyanobacteria, as shown here, are complexed to bioavailable, polar glycolipids. Human and rodent studies show that high doses of echium oil reduce circulating triglycerides (15, 67) and that >1 g/day of SDA from transgenic soybean oil is effective at raising tissue membrane levels of EPA and improving the omega-3 index in humans (erythrocyte EPA and DHA) (21, 68-70). SDA-enriched soybean oil fed to laying hens performs better than ALA at enriching eggs with n-3 LC-PUFAs and particularly EPA (17), and similar effects were obtained with meat from broiler chickens (71). Echium oil also enhances total n-3 PUFA levels, including EPA, in the milk of dairy cattle (72). However, there continues to be substantial barriers to the supplementation of SDA complexed to triglycerides (as found in seed and soybean oils), and this has limited the widespread use of SDA; furthermore, initial studies suggest that PUFAs or LC-PUFAs complexed to MGDG and DGDG may provide greater bioavailability than non-polar, triglyceride-containing oils and phospholipids found in krill oil (31).

## Conclusions

Here we demonstrate that cyanobacteria, and especially *Leptolyngbya* BL0902, bioengineered with cyanobacterial lipid biosynthetic promoting genes, produce large quantities of SDA up to about 27% of total fatty acids. Importantly, both newly-synthesized SDA and ETA are found conjugated to galactose-containing glycerol backbones in what initial studies indicate is a more bioavailable polar lipid form than the neutral storage lipids like triacylglycerols of oils. Additionally, several novel and potentially beneficial molecular species of MGDG and DGDG are formed in these cyanobacteria that may serve as highly bioavailable, anti-inflammatory compounds. SDA-producing cyanobacteria as developed here are thus promising, sustainable sources of omega-3 PUFAs that could replace unstable fish oil products (73, 74) and fish meal as nutritional supplements for human, agricultural and aquacultural use.

## Methods

### Molecular biology approaches

Overall, the goal of the project was to generate, for testing in various cyanobacteria, a set of plasmids expressing one, two or three cyanobacterial genes that we hypothesized would enable stearidonic acid (SDA) production in a wide range of cyanobacteria (as long as they were already able to produce the dienoic fatty acid linoleic acid, 18:2). The three genes of interest for this work were *des*D, *des*B and *vipp*1, for which we designed coding sequences based on the corresponding protein sequences from well documented sources: DesD, the acyl-lipid Δ6-desaturase, was from *Synechocystis* sp. PCC 6803; DesB, the acyl-lipid ω3-desaturase (alternatively named methyl-end or Δ15 desaturase) was from *Synechococcus* sp. PCC 7002; and Vipp1, dubbed the “Vesicle-inducing protein in plastids,” was from *Synechococcus* sp. PCC 7002. Each protein sequence was used to design the synthetic gene, codon optimized for expression in *E. coli* (given a lack of options for cyanobacteria). Each cloned insert obtained from GenScript was then amplified by polymerase chain reaction (PCR) and cloned into the Gateway donor plasmid, pENTR/SD/D-topo (Invitrogen), which provides an upstream Shine-Dalgarno sequence (ribosome-binding site) that is known to function in cyanobacteria. Sequences of all plasmid inserts were verified and transferred into pAM4418 using the Gateway recombination system (Invitrogen) and Invitrogen’s LR Clonase II Enzyme Mix, with verification of positive clones by restriction digestion. Details of the cloning steps varied with the construct and are given below. The expression vector used, pAM4418, is a broad host range, *E. coli*-cyanobacteria shuttle plasmid that confers resistance to streptomycin and spectinomycin and contains both the lacI^q^ repressor and the *trc* promoter from *E*.*coli*, in addition to the Gateway recombination cassette (44).

Using the overall approaches outlined above, the following seven expression plasmids were generated: (1) pD: encodes DesD, the Δ6 desaturase; (2) pB: encodes DesB, the Δ15 desaturase; (3) pDB: encodes both DesD and DesB; (4) pV: encodes Vipp1; (5) pDV: encodes both DesD and Vipp1; (6) pBV: encodes both DesB and Vipp1; and (7) pDBV: encodes all three, DesD, DesB and Vipp1.. The contents of these vectors are also summarized at the bottom of Fig. 2B.

In order to create the engineered pENTR plasmids to generate pD and pB, *des*D and *des*B, respectively, were PCR amplified using primers which added the sequence CACC before the initiating ATG codon and a *Xho*1 restriction site following the termination codon, enabling directional cloning of the PCR product into pENTR/SD/D-topo. For the next two constructs, pDV and pBV, the downstream *Xho*I restriction sites after each desaturase-encoding gene, in combination with the *Asc*I site in the pENTR/SD/D-topo plasmid, provided sites for insertion of *vipp*1; the *vipp*1-containing fragment was excised from the Genscript plasmid using *Xho*I and *Asc*1, then ligated following each of the two desaturase genes into the pENTR/SD/D-topo clones described above. To create the pENTR plasmid encoding only Vipp1 (to generate pV), *vipp*1 from Genscript was PCR amplified as described above for *des*D and *des*B to enable directional cloning of the PCR product into pENTR/SD/D-topo.

In order to create pDBV, the *desB* sequence from GenScript was amplified by PCR to introduce a *Hin*dIII site plus a ribosome-binding site on the 5’ end and an *Asc*1 restriction site on the 3’ end. The PCR product was digested with *Hin*dIII and *Asc*I and ligated into the pENTR/SD/D-topo plasmid already containing *des*D and *vipp*1, digested with the same two restriction enzymes.

Finally, the pENTR plasmid used to generate pDB, encoding only DesD and DesB, was derived from the pENTR plasmid containing all three genes (above) by digestion with *Hin*dIII and *Xho*I to remove the Vipp1-encoding gene, filling in the ends with dNTPs and DNA polymerase (Klenow fragment), then ligating the blunt ends together.

### Genetic engineering, growth and harvest of cyanobacterial strains

The seven pAM4418-derived plasmids were used to transform *E*.*coli* DH10B cells containing the conjugal and helper plasmids, pRL443 and pRL623, respectively. Transformants were grown overnight in rich LB media, washed with fresh LB, and resuspended in BG-11 media as a 10-fold concentrated stock. Cultures of the three host cyanobacteria (*Leptolyngbya* sp. strain BL0902, *Anabaena* sp. PCC7120 in BG-11 media and *Synechococcus* sp. PCC7002 in Medium A (75)) were grown to late exponential phase, harvested by centrifugation and washed twice with fresh media, before resuspension as a 4-fold concentrated stock. Cyanobacterial suspensions were sonicated in a bath for 10 min to reduce the length of the multicellular strands, then mixed with DH10B transformants. Cell mixtures were centrifuged, resuspended in 200 µL of BG-11 media, incubated for 1 h at 30 °C, then spread on BG-11/5 % LB agar plates. After incubation for 24 h in low light at 30°C, cells were washed and spread on BG-11 agar containing 2 µg/mL spectinomycin and streptomycin. After 7-10 days incubation at 30°C under illumination [∼20-30 µmol photons/(m^2^ · s)], single colonies were restreaked onto a fresh antibiotic-containing plate and incubated for 5-7 days. Bacteria scraped from the plate were transferred to 30 mL of BG-11 in a 250 mL conical flask, grown for 5 days at 30 °C (with shaking at 120 rpm and illumination), then harvested by centrifugation (5,000 rpm for 10 min). For *Synechococcus* sp. PCC7002, Medium A replaced BG-11 media in all steps of the conjugation and growth. Cell pellets were stored frozen at -80 °C, then dried by lyophilization and weighed before the analysis of lipid content.

### Characterization of the Fatty Acid Content of Cyanobacteria Lipids

Lyophilized cell pellets were extracted utilizing a modified Bligh/Dyer for total fatty acid analysis (∼2 mg/sample). For total fatty acid analysis, solvents were evaporated under a stream of nitrogen in the presence of a fatty acid internal standard (triheptodecanoin, 10 µg). The dried extract was then subjected to base hydrolysis and derivatization in the presence of boron trifluoride (5 min, 100°C) to form fatty acid methyl esters (FAME) following a modification of the protocol by Metcalfe *et al*. (76) as previously described (77, 78). FAMEs were analyzed on an Agilent J&W DB-23 column (30 m × 0.25 mm ID, film thickness 0.25 μm) using HP 5890 gas chromatography (GC) with a flame ionization detector (FID). Individual fatty acids were identified by their elution times relative to authenticated fatty acid standards, and fatty acid quantities were determined by their abundance relative to the internal standard.

For lipidomics analysis, total lipid extracts derived from 2 mg lyophilized biomass from wild type and pDBV-modified strains of *Leptolyngbya* sp. strain BL0902 and *Synechococcus* sp. PCC7002 were dried as above and dissolved in 100 µL isopropyl alcohol/methanol (50:50) for LC-MS/MS analysis. Samples (10% of total per injection) were analyzed on a high resolution Q Exactive HF Hybrid Quadrupole - Orbitrap Mass Spectrometer equipped with a heated electrospray ionization (HESI)-II source (Thermo Scientific, Rockford, IL) and a Vanquish Horizon UHPLC system (Thermo Scientific, Rockford, IL), with source parameters as follows: sheath gas flow rate, 40 L/min; auxiliary gas flow rate, 5 L/min; spray voltage, 4.0 kV with positive mode and 3.5 kV with negative mode; capillary temperature, 350°C; S-lens RF voltage, 75 V.

Chromatographic separation was achieved on an Accucore C30 column (2.6µm, 3mm x 150mm, Thermo Scientific, Rockford, IL) with linear gradient elution consisting of mobile phases A (water/acetonitrile = 40:60) and B (isopropyl alcohol/acetonitrile = 90:10) at 0.35 mL/min. Both mobile phases contained 0.1% formic acid and 10 mM ammonium formate and the gradient was from 40% B at 0 min to 95% B at 30 min.

MS spectra were acquired by data-dependent scans in positive and negative mode. A survey scan was performed at MS1 level to identify top ten most abundant precursor ions followed by MS2 scans where product ions were generated from selected ions. High-energy collisional dissociation (HCD) was utilized for ion fragmentation with stepped collision energy of 25/30 eV and 30/50/100 eV in each positive and negative polarity (79). The dynamic exclusion option was enabled during data-dependent scans to enhance compound identification in complex mixtures. Acquired spectra were processed using LipidSearch software v4.1 (Thermo Scientific, Rockford, IL) with the selection of following classes of lipids: lysophosphatidylcholine (LPC), phosphatidylcholine (PC), lysophosphatidylethanolamine (LPE), phosphatidyl-ethanolamine (PE), lysophosphatidylserine (LPS), phosphatidylserine (PS), lysophosphatidylglycerol (LPG), phosphatidylglycerol (PG), lysophosphatidylinositol (LPI), phosphatidylinositol (PI), lysophosphatidic acid (LPA), phosphatidic acid (PA), sphingomyelin (SM), phytosphingosine (phSM), monoglyceride (MG), diglyceride (DG), triglyceride (TG), fatty acid (FA), (O-acyl)-1-hydroxy fatty acid (OAHFA), cardiolipin (CL), sphingoshine (So), sphingoshine phosphate (SoP), glucosylsphingoshine (SoG1), monoglycosylceramide (CerG1), diglycosylceramide (CerG2), triglycosylceramide (CerG3), ceramides (Cer), monosialotetrahexosylganglioside (GM2), cholesteryl ester (ChE), zymosteryl (ZyE), stigmasteryl ester (StE), sitosteryl ester (SiE), coenzymes (Co), monogalactosylmonoacylglycerol (MGMG), monogalactosyl-diacylglycerol (MGDG), digalactosylmonoacylglycerol (DGMG), digalactosyldiacylglycerol (DGDG), sulfoquinovosylmonoacylglycerol (SQMG), and sulfoquinovosyldiacylglycerol (SQDG). Parameters for the product search workflow were: precursor mass tolerance, 5 ppm; product mass tolerance, 5 ppm; product ion intensity threshold, 1.0% relative to precursor; matching score threshold, 2.0. All peak areas were normalized to the total ion current (the total area under the curve in the chromatogram).

## Abbreviations

LC: Long chain
FA: fatty acid
PUFA: polyunsaturated fatty acid; *des*B and DesB, Δ15 acyl lipid desaturase (gene and protein, respectively); *des*D and DesD Δ6 acyl lipid desaturase (gene and protein, respectively)
Vipp1: Vesicle-inducing protein in plastids 1
DesA: Δ12 acyl lipid desaturase
DesC: Δ9 acyl lipid desaturase
SDA: stearidonic acid (18:4, 18 carbons with 4 double bonds, positions 6, 9, 12, 15)
ALA: alpha-linolenic acid (18:3, double bond positions 9, 12, 15)
ETA: eicosatetraenoic acid (20:4, double bond positions 8, 11, 14, 17)
GLA: gamma-linolenic acid (18:3, double bond positions 6, 9, 12)
LA: linoleic acid (18:2, double bond positions 9, 12)
MGDG: monogalactosyldiacylglycerol
DGDG: digalactosyl-diacylglycerol
SQDG: sulfoquinovosyldiacylglycerol
PG: phosphatidyl glycerol
pDBV: plasmid derived from pAM4418 containing genes encoding DesD, DesB and Vipp1
EPA: eicosapentaenoic acid (20:5, double bond positions 5, 8, 11, 14, 17)
DHA: docosahexaenoic acid (22:6, double bond positions 4, 7, 10, 13, 16, 19)
CoA: coenzyme A
ACP: acyl carrier protein
BL0902: *Leptolyngbya* sp. strain BL0902
7002: *Synechococcus* sp. PCC7002
7120: *Anabaena* sp. PCC7120
GC-FID: gas chromatography-flame ionization detection
COX1 and COX2: cyclooxygenases 1 and 2
PCR: polymerase chain reaction
FAME: fatty acid methyl esters
LC-MS/MS: liquid chromatography-tandem mass spectrometry
UHPLC: ultrahigh performance liquid chromatography.

## Acknowledgements

We thank Wake Forest Innovations for financial support through the Value Inflection, Commercialization Pathway and Catalyst Awards, and particularly Sarah Haigh Molina for her efforts in supporting this work. We also acknowledge contributions from the National Institutes of Health, R01 AT008621 (to F.H.C.) and R01 GM119227 and R35 GM135179 (to L.B.P.). We are grateful to Dr. James Golden (University of California at San Diego) for providing the pAM4418 plasmid and the strains of cyanobacteria tested in this work, as well as advice regarding growth of cyanobacteria. We thank Deena Wykle for assistance with literature searches. We also acknowledge the support provided by the Proteomics and Metabolomics Shared Resource of the Wake Forest Baptist Comprehensive Cancer Center (NIH/NCI P30 CA12197; facility Director Dr. Cristina M. Furdui).

