## Supplemental data for "Acyl-lipid desaturases and Vipp1 cooperate in cyanobacteria to produce novel omega-3 PUFA-containing glycolipids"

### Supplementary Material

Included below are additional results related to Bioinformatics, total fatty acid contents in all three strains of cyanobacteria studied (Fig. S1), contributions of individual genes to the PUFA-promoting effect of pDBV (Fig. S2), lipidomics data from the second experiment (Table S1) and the identity of the second acyl chain in n-3 PUFA-containing lipids (Fig. S3), summary of ALA and SDA production by pBV and pDBV-containing *Leptolyngbya* sp. strain BL0902 (Table S2), and sample MS2 spectra used for peak identification (Fig. S4). Two additional Excel spreadsheets of supplementary data are also included:

Supplementary Excel Data. Fatty acid profile for cyanobacteria with and without plasmid constructs (Excel)

#### Results

**Bioinformatics to search for *des*B and *des*D loci in *Leptolyngbya* species.** To complement our fatty acid analyses of wild type *Leptolyngbya* sp. strain BL0902 for which there is not yet a published sequence, we conducted searches for the well conserved Δ6 (*des*D) and ω3 (*des*B) desaturase genes based on the known sequences from *Synechocystis* sp. PCC6803, as well as variants from several

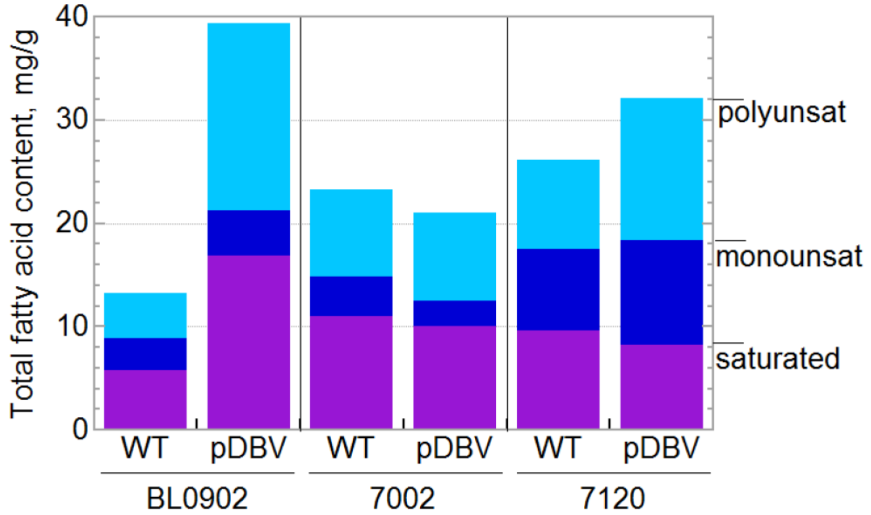


**Figure S1.** Total fatty acid contents of wild type and engineered cyanobacterial strains. Three cyanobacterial strains, *Leptolyngbya* sp. strain BL0902, *Synechococcus* sp. PCC 7002 and *Anabaena* sp. PCC 7120, were selected for testing given their favorable starting contents of linoleic acid (18:2). Wild type (WT) or engineered (+pDBV) strains were grown, harvested and analyzed for fatty acid content (FAME analysis) as described in Fig. 1 and Methods. pDBV includes three cyanobacterial genes encoding DesD (Δ6 desaturase), DesB (omega-3 or Δ15 desaturase), and the "vesicle-inducing protein in plastids" (Vipp1). Shown are total saturated (magenta), monounsaturated (dark blue) and polyunsaturated (cyan) fatty acid contents (n=5, 3, 3, 3, 2 and 1 from left to right).

other cyanobacterial species to confirm the results. A Blastp search for *des*B in *Leptolyngbya* species at the National Center for Biomedical Information (https://blast.ncbi.nlm.nih.gov/) in March 2020 returned 22 very convincing homologues (e-value < 2E-168) using the non-redundant protein sequence database (nr), whereas no convincing *des*D ortholog was identified in a parallel search (e-values all > 1E-14).

**Supplemental data reporting cyanobacterial fatty acids and lipids.** Reported below are total fatty acid contents in all three strains of cyanobacteria studied (Fig. S1), contributions of individual genes to the PUFA-promoting effect of pDBV (Fig. S2), and the identity of the second acyl chain in n-3 PUFA-containing lipids based on the lipidomics analysis (Fig. S3).

**Lipidomics of *Leptolyngbya* sp. strain BL0902 and *Synechococcus* sp. PCC7002.** Analyses were conducted as described in the main manuscript, with wild type and pDBV-containing forms of each of two cyanobacterial strains (n=1 each). In this second experiment, ALA, SDA and ETA were found in a larger number of lipid species, and SQDG species were more enriched in SDA and particularly ALA in this experiment, whereas PG rather than SQDG contained measureable amounts of these fatty acids in the first experiment (Table S1).

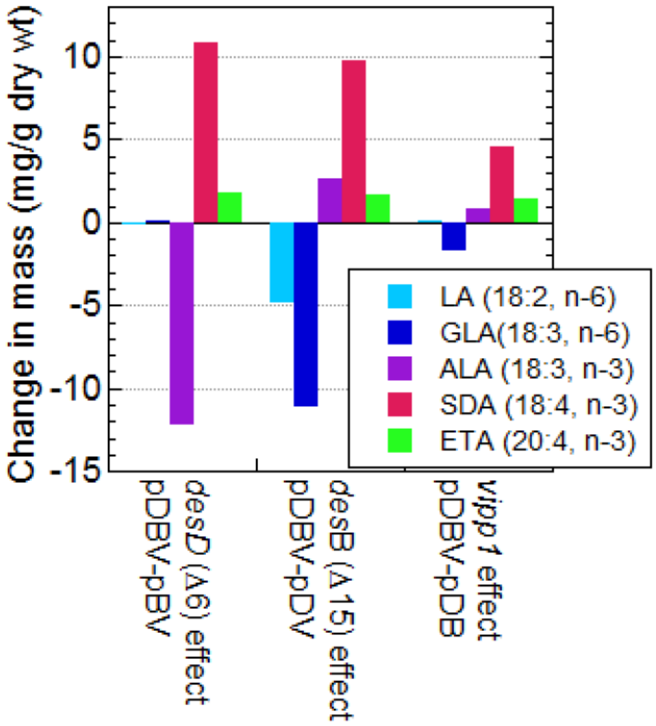


**Figure S2.** Individual contribution toward the PUFA profile (generated by pDBV addition) of the engineered cyanobacterial genes in the context of the presence of the other two genes. Based on FAME analysis of the PUFAs depicted in Figure 3, changes in mass of each of the 18C- and 20C-PUFAs are shown, with conversions observed consistent with each of the expected molecular functions.

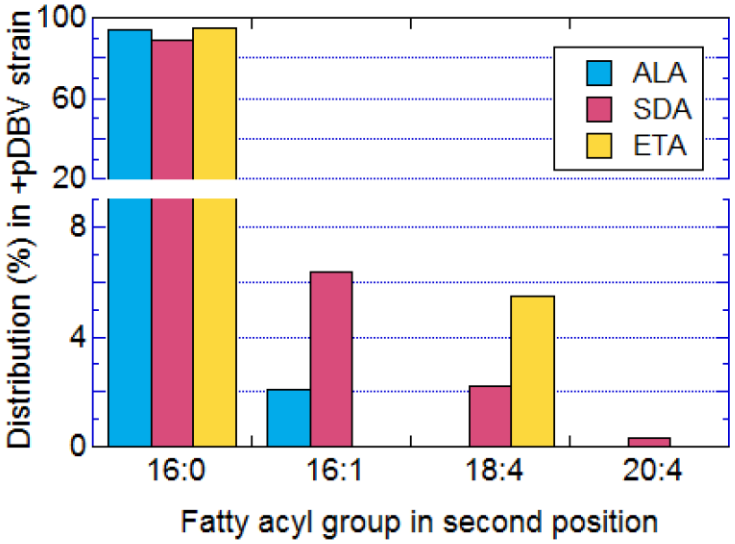


**Figure S3.** Selective distribution of major n-3 PUFAs (ALA, SDA and ETA) within *Leptolyngbya* BL0902 glycolipids with respect to the nature of the second of the two acyl chains within each lipid molecule, based on lipidomics analyses depicted in Figure 4.

**Table S1.** Molecular species of lipids by LC-MS/MS in wild type (WT) and engineered (+pDBV) *Leptolyngbya* sp. strain BL0902 and *Synechococcus* sp. PCC7002 (n=1 each).[a]

| Molecular species | Cyano strain | MGDG | | DGDG | | SQDG | | PG | |
|---|---|---|---|---|---|---|---|---|---|
| | | WT | +pDBV | WT | +pDBV | WT | +pDBV | WT | +pDBV |
| 14:0/16:0 | BL0902 | 0.111 | 0.850 | 0.000 | 0.000 | | | | |
| | 7002 | 0.225 | 0.112 | 0.012 | 0.008 | | | | |
| 14:0/18:3 | BL0902 | 0.008 | 0.042 | 0.012 | 0.008 | | | 0.000 | 0.009 |
| | 7002 | 0.017 | 0.008 | 0.015 | 0.001 | | | 0.001 | 0.010 |
| 14:0/18:4 | BL0902 | 0.000 | 0.168 | 0.001 | 0.012 | | | | |
| | 7002 | 0.000 | 0.022 | 0.003 | 0.004 | | | | |
| 16:0/16:0 | BL0902 | 0.051 | 0.796 | 0.000 | 0.012 | 0.005 | 0.006 | 0.001 | 0.256 |
| | 7002 | 0.053 | 0.030 | 0.000 | 0.004 | 0.027 | 0.008 | 0.015 | 0.032 |
| 16:0/16:1 | BL0902 | 0.090 | 0.508 | 0.001 | 0.045 | | | 0.003 | 0.105 |
| | 7002 | 0.072 | 0.022 | 0.002 | 0.004 | | | 0.018 | 0.039 |
| 16:0/16:3 | BL0902 | 0.000 | 0.051 | | | | | | |
| | 7002 | 0.000 | 0.014 | | | | | | |
| 16:0/17:2 | BL0902 | 0.019 | 0.065 | | | | | 0.000 | 0.010 |
| | 7002 | 0.057 | 0.030 | | | | | 0.011 | 0.005 |
| 16:0/17:4 | BL0902 | 0.000 | 0.024 | | | | | | |
| | 7002 | 0.000 | 0.015 | | | | | | |
| 16:0/18:1 | BL0902 | 0.629 | 1.479 | 0.000 | 0.000 | | | 0.207 | 1.164 |
| | 7002 | 0.676 | 0.273 | 0.005 | 0.003 | | | 0.640 | 0.353 |
| 16:0/18:2 | BL0902 | 0.566 | 3.214 | 0.011 | 0.134 | | | 0.057 | 1.843 |
| | 7002 | 3.964 | 0.979 | 0.030 | 0.056 | | | 1.710 | 0.266 |
| 16:0/18:3 | BL0902 | 0.168 | 1.582 | 0.017 | 0.058 | 0.128 | 2.301 | | |
| | 7002 | 0.692 | 0.858 | 0.325 | 0.011 | 0.796 | 4.914 | | |
| 16:0/18:4 | BL0902 | 0.001 | 5.330 | 0.012 | 0.212 | 0.004 | 1.109 | 0.000 | 0.035 |
| | 7002 | 0.001 | 2.199 | 0.126 | 0.123 | 0.000 | 0.235 | 0.000 | 0.013 |
| 16:0/18:5 | BL0902 | 0.001 | 0.009 | | | | | | |
| | 7002 | 0.025 | 0.008 | | | | | | |
| 16:0/19:2 | BL0902 | 0.007 | 0.011 | | | | | | |
| | 7002 | 0.029 | 0.005 | | | | | | |
| 16:0/20:4 | BL0902 | 0.053 | 0.527 | 0.000 | 0.712 | | | 0.000 | 0.114 |
| | 7002 | 0.015 | 0.145 | 0.000 | 0.295 | | | 0.000 | 0.035 |
| 16:0/20:5 | BL0902 | | | 0.006 | 0.038 | 0.031 | 0.023 | | |
| | 7002 | | | 0.003 | 0.011 | 0.019 | 0.023 | | |
| 16:1/18:2 | BL0902 | 0.483 | 0.122 | 0.000 | 0.000 | | | 0.010 | 2.403 |
| | 7002 | 0.580 | 0.021 | 0.003 | 0.000 | | | 0.464 | 2.173 |

| | | | | | | | | | |
|---|---|---|---|---|---|---|---|---|---|
| 16:1/18:3 | BL0902 | 0.100 | 0.180 | 0.048 | 0.014 | 0.014 | 0.088 | 0.017 | 0.046 |
| | 7002 | 0.161 | 0.163 | 0.004 | 0.004 | 0.001 | 0.059 | 0.018 | 0.057 |
| 16:1/18:4 | BL0902 | 0.001 | 1.006 | 0.013 | 0.467 | | | | |
| | 7002 | 0.000 | 0.363 | 0.116 | 0.282 | | | | |
| 16:2/18:3 | BL0902 | | | 0.000 | 0.025 | | | | |
| | 7002 | | | 0.000 | 0.010 | | | | |
| 16:2/18:4 | BL0902 | 0.013 | 0.117 | 0.000 | 0.014 | | | | |
| | 7002 | 0.003 | 0.053 | 0.005 | 0.008 | | | | |
| 16:3/18:4 | BL0902 | 0.000 | 0.021 | 0.000 | 0.053 | | | | |
| | 7002 | 0.000 | 0.011 | 0.000 | 0.033 | | | | |
| 17:1/18:2 | BL0902 | 0.019 | 0.003 | | | | | | |
| | 7002 | 0.051 | 0.003 | | | | | | |
| 17:1/18:4 | BL0902 | 0.000 | 0.016 | | | | | | |
| | 7002 | 0.000 | 0.017 | | | | | | |
| 17:2/18:4 | BL0902 | 0.000 | 0.014 | 0.000 | 0.000 | | | | |
| | 7002 | 0.000 | 0.011 | 0.003 | 0.002 | | | | |
| 17:3/18:4 | BL0902 | 0.000 | 0.007 | 0.000 | 0.003 | | | | |
| | 7002 | 0.000 | 0.008 | 0.000 | 0.004 | | | | |
| 18:0/18:4 | BL0902 | 0.005 | 0.012 | | | | | | |
| | 7002 | 0.001 | 0.011 | | | | | | |
| 18:1/18:1 | BL0902 | 0.199 | 0.030 | 0.000 | 0.000 | 0.090 | 0.028 | 0.018 | 0.306 |
| | 7002 | 0.081 | 0.013 | 0.004 | 0.001 | 0.134 | 0.030 | 0.070 | 0.014 |
| 18:1/18:2 | BL0902 | 0.479 | 0.114 | 0.007 | 0.000 | | | 0.015 | 0.050 |
| | 7002 | 0.272 | 0.070 | 0.005 | 0.000 | | | 0.041 | 0.005 |
| 18:1/18:4 | BL0902 | 0.194 | 0.470 | | | | | | |
| | 7002 | 0.222 | 0.256 | | | | | | |
| 18:1/18:5 | BL0902 | | | | | 0.003 | 0.065 | | |
| | 7002 | | | | | 0.002 | 0.065 | | |
| 18:1/20:4 | BL0902 | 0.000 | 0.014 | | | | | | |
| | 7002 | 0.002 | 0.007 | | | | | | |
| 18:2/18:2 | BL0902 | 0.330 | 0.027 | 0.029 | 0.000 | | | 0.003 | 0.017 |
| | 7002 | 0.554 | 0.041 | 0.032 | 0.001 | | | 0.008 | 0.017 |
| 18:2/18:3 | BL0902 | 0.079 | 0.207 | 0.028 | 0.019 | | | | |
| | 7002 | 0.281 | 0.106 | 0.186 | 0.013 | | | | |
| 18:2/18:4 | BL0902 | 0.005 | 0.097 | 0.011 | 0.026 | | | | |
| | 7002 | 0.001 | 0.022 | 0.127 | 0.031 | | | | |
| 18:2/18:5 | BL0902 | | | | | 0.000 | 0.088 | | |
| | 7002 | | | | | 0.000 | 0.031 | | |

| | | | | | | | | | |
|---|---|---|---|---|---|---|---|---|---|
| 18:2/20:4 | BL0902 | 0.000 | 0.019 | | | | | | |
| | 7002 | 0.001 | 0.015 | | | | | | |
| 18:3/18:3 | BL0902 | 0.013 | 0.405 | 0.000 | 0.020 | | | 0.000 | 0.001 |
| | 7002 | 0.030 | 0.094 | 0.000 | 0.014 | | | 0.001 | 0.026 |
| 18:3/18:4 | BL0902 | 0.000 | 0.502 | 0.010 | 0.087 | | | | |
| | 7002 | 0.001 | 0.261 | 0.015 | 0.063 | | | | |
| 18:3/20:3 | BL0902 | 0.000 | 0.006 | | | | | | |
| | 7002 | 0.000 | 0.009 | | | | | | |
| 18:3/20:4 | BL0902 | 0.000 | 0.029 | 0.000 | 0.210 | | | | |
| | 7002 | 0.000 | 0.016 | 0.000 | 0.111 | | | | |
| 18:4/18:4 | BL0902 | 0.000 | 0.868 | 0.000 | 0.014 | 0.000 | 0.075 | | |
| | 7002 | 0.000 | 0.203 | 0.000 | 0.013 | 0.000 | 0.006 | | |
| 18:4/20:4 | BL0902 | 0.000 | 0.114 | 0.000 | 0.167 | | | | |
| | 7002 | 0.000 | 0.048 | 0.000 | 0.049 | | | | |
| 18:4/20:5 | BL0902 | 0.003 | 0.020 | | | | | | |
| | 7002 | 0.002 | 0.014 | | | | | | |

[a]Values reported are estimates of mg/g of total fatty acid based on (i) normalized peak areas from LC/MS, (ii) fraction of total peak area for each species in a sample, and (iii) known total fatty acid yield for that organism from GC-FID analysis. This treatment assumes that all species exhibit the same ionization efficiency. Shown are mean +/- standard deviation for wild type (WT) *Leptolyngbya* BL0902 without and with pDBV, and *Synechococcus* sp. PCC 7002 (n=1 each). No entry means that the species was not observed in the WT or pDBV samples in that category.

[b]Molecular species shown are for the two fatty acyl chains, giving carbon chain length and number of double bonds for each.

[c]MGDG = monogalactosyldiacylglycerol; DGDG = digalactosyldiacylglycerol; SQDG = sulfoquinovosyldiacylglycerol; PG = phosphatidyl glycerol

**Table S2.** Content (reporting mass and mol% of total fatty acids) of 18-carbon omega3 fatty acids in engineered *Leptolyngbya* sp. strain BL0902.

| Species | | ALA (18:3, n-3), mg/g and mol% of total FAs[a] | SDA (18:4, n-3) mg/g and mol% of total FAs[a] |
|---|---|---|---|
| *Leptolyngbya* sp. strain BL0902 | None (wt) | 3.1 ± 1.6 mg/g<br>22.6 ± 11.5 % | n.d.[b] |
| | With pBV (to express DesB and Vipp1)[c] | 16.0 ± 0.7 mg/g<br>37.7 ± 1.7 % | n.d. |
| | With pDBV (to express DesD, DesB and Vipp1)[c] | 3.8 ± 0.4 mg/g<br>9.3 ± 0.9 % | 10.8 ± 0.4 mg/g<br>26.6 ± 1.0% |

[a] Expressed as mean ± standard deviation.

[b] n.d. = none detected

[c] pBV encodes *Synechococcus* sp. PCC7002 DesB for Δ15 desaturase (also known as ω3 or methyl end desaturase) and *Synechococcus* sp. PCC7002 Vipp1 for inducing thylakoid membranes; pDBV is the three-gene plasmid derived from pAM4418 (Fig. 2) developed in this work which includes the two genes of pBV plus *Synechocystis* sp. PCC6803 *des*D for Δ6 desaturase expression.

**Accumulation of ALA in *Leptolyngbya* BL0902 with pBV.** Although our goal in this study was to achieve significant SDA production by cyanobacteria, ALA is also a potentially important omega-3 fatty acid with greater stability that EPA and DHA. ALA accumulated in these studies only in the absence of *des*D (the Δ6 desaturase); our pBV plasmid enabled significant levels of up to 37.7% to accumulate. Similar experiments by others at relatively low illumination like ours [<u><</u>50 µmol photons/($m^2 \cdot s$)] achieved about 23 to 28% ALA accumulation, an amount that is similar to that present in wild type *Leptolyngbya* BL0902 (Table 3). Yoshino et al. (1) reported accumulation of as much as 53% ALA under conditions of high incident light [150 µmol photons/($m^2 \cdot s$)], but in those experiments (which included *des*D overexpression), SDA reached a maximum of only 6.2%.

**Confirmation of mass spectrometry peak identification through sample MS-MS spectra, and combined approaches to track SDA distribution among lipid species.** As an illustration of the quality of the MS-MS data used for identification of lipid species in the cyanobacterial samples, two MS2 spectra of key species of interest (but at relatively low abundance) are shown along with labeling of several of the major peaks that provide the information for identification (Fig. S4).

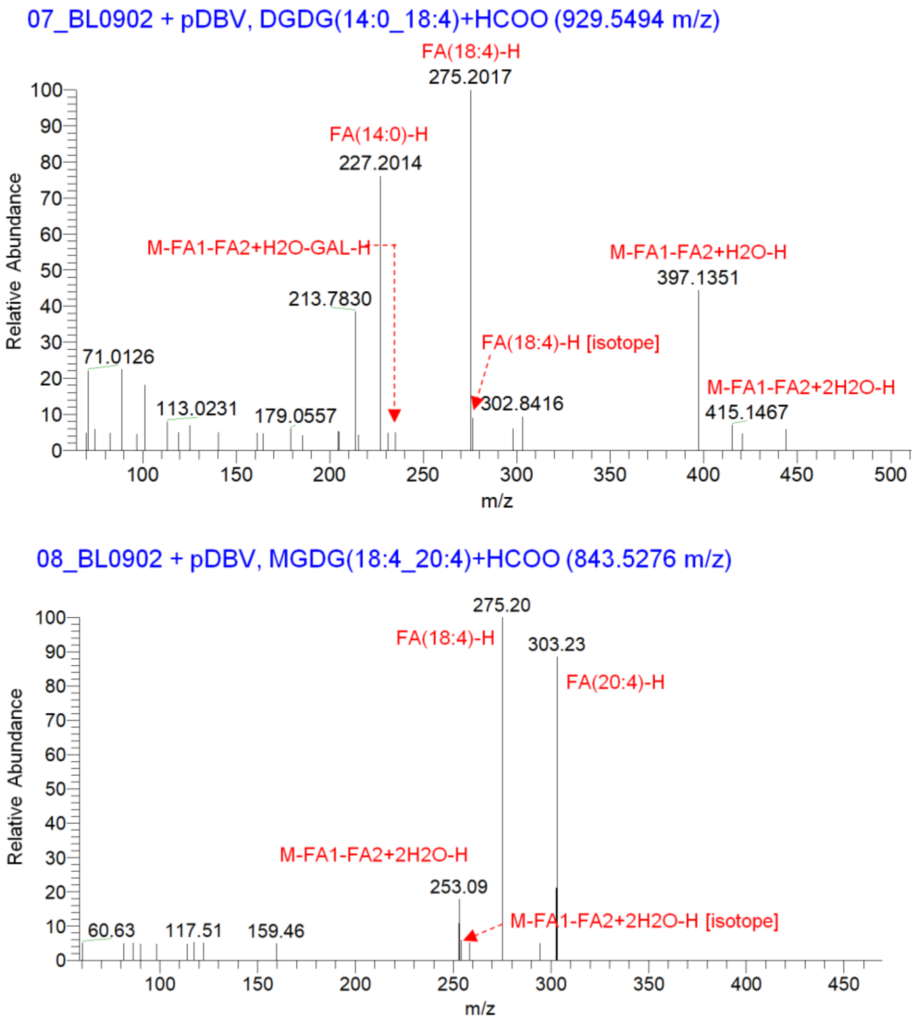


**Figure S4.** Spectra from MS-MS of two lipid species of interest. Shown are the fragment ions and identifying features for lipids from two samples of *Leptolyngbya* sp. strain BL0902 with pDBV. The upper panel shows DGDG with 14:0 and 18:4 fatty acyl chains (parent ion 929.5494 m/z), and the lower panel shows MGDG with 18:4 and 20:4 acyl chains (parent ion 843.5276 m/z). All fragments labeled in red are perfect matches.
